# Emergence of *Candida auris* in Minas Gerais, Brazil: Genomic Surveillance to inform Rapid Public Health Responses

**DOI:** 10.1101/2024.12.13.627932

**Authors:** Luiz Marcelo Ribeiro Tomé, Dhian Renato Almeida Camargo, Rafael Wesley Bastos, Sara Cândida Ferreira dos Santos, Natália Rocha Guimarães, Sílvia Helena Sousa Pietra Pedroso, Paulo Eduardo de Souza da Silva, Aristóteles Góes-Neto, Lida Jouca de Assis Figueredo, Gabriella da Côrte Castro, Ana Maria Ribeiro Nunes Rodrigues, Flavia Ribeiro Soares Cruzeiro, Nádia Aparecida Campos Dutra, Josiane Barbosa Piedade Moura, Glauco de Carvalho Pereira, Carmem Dolores Faria, Marta Giovanetti, Felipe Campos de Melo Iani, Luiz Carlos Júnior Alcantara, Talita Émile Ribeiro Adelino

**Affiliations:** Instituto René Rachou – Fiocruz Minas, Belo Horizonte, Minas Gerais, Brazil; Central Public Health Laboratory of Minas Gerais (Lacen-MG), Ezequiel Dias Foundation – Funed, Belo Horizonte, Minas Gerais, Brazil; Universidade Federal do Rio Grande do Norte – UFRN, Natal, Rio Grande do Norte, Brazil; Universidade Federal de Minas Gerais – UFMG; Secretaria de Estado de Saúde de Minas Gerais – SES-MG, Belo Horizonte, Minas Gerais, Brazil; Universita Campus Bio-Medico di Roma, Italy; CT Vacinas – Universidade Federal de Minas Gerais, Belo Horizonte, Minas Gerais, Brazil

## Abstract

*Candida auris* poses a significant global health threat due to its multidrug-resistance and outbreak potential. In this study, we report its emergence in Minas Gerais, Brazil, supported by genomic surveillance that identified Clade-IV isolates, suggested a potential introduction from Colombia, and detected a missense mutation associated with azole resistance.

## Introduction

*Candida auris* (*Candidozyma auris*) is an emerging multidrug-resistant (MDR) fungal pathogen that poses a significant challenge to global public health. Since its initial identification in Japan in 2009, *C. auris* has spread worldwide, raising substantial concern due to its resistance to antifungal treatments, high transmissibility, and persistence in healthcare environments (1). Despite its clinical importance, the evolutionary origins of *C. auris* remain poorly understood, although they are speculated to be linked to climate change and the increasing population of immunocompromised individuals (2).

Circulating strains of *C. auris* have been classified into six genomic clades, named according to their geographic region of origin: Clade-I (South Asian), Clade-II (East Asian), Clade-III (African), Clade-IV (South American), Clade-V (Iranian), and Clade-VI (Singaporean) (3,4). This classification is based on genomic divergence, assessed through single nucleotide polymorphism (SNP) analyses, antifungal susceptibility profiles, and outbreak potential (3).

In Brazil, *C. auris* was first detected in December 2020 in Salvador, Bahia (5). The isolate, belonging to Clade-I, was susceptible to key antifungals, including amphotericin B, fluconazole, voriconazole, and anidulafungin (6). By late 2021, *C. auris* from Clade-IV was identified in Pernambuco, with isolates showing sensitivity to amphotericin B, azoles, and echinocandins (7). Since then, Pernambuco has reported over five outbreaks and more than 48 cases, including colonization and infections. In June 2023, a positive case involving a newborn was reported in São Paulo (ANVISA, 2023).

In September 2024, *C. auris* was detected for the first time in a hospital in Belo Horizonte, Minas Gerais, with four positive cases identified. Preliminary evidence points to a possible introduction from Colombia. In response to the emergence of this MDR pathogen in Minas Gerais and its outbreak potential, we promptly characterized the isolates and implemented fungal genomic surveillance at the Central Public Health Laboratory of Minas Gerais (Lacen-MG), located at Ezequiel Dias Foundation (Funed).

### The Study

Clinical samples (urine) and swabs from the axillary and inguinal regions of patients admitted to the Intensive Care Unit (ICU) with suspected *C. auris* infection or colonization were collected at a hospital in Belo Horizonte (Table 1). Additionally, environmental swabs were obtained from ICU devices. This study was approved by the research ethics committee of the Ezequiel Dias Foundation (CAAE: 85347524.5.0000.9507).

**Table 1.** Demographic data of patients and details of *Candida auris* positive samples sequenced in this study.

| Patient | Sample ID | State | Municipality | Sample Source | Biological Material | Paciente Age (years) | Sex | Collection date |
| --- | --- | --- | --- | --- | --- | --- | --- | --- |
| 1 | FUNED538* | Minas Gerais | Belo Horizonte | Clinical Sample | Urine | 58 | Male | 2024-09-17 |
| 2 | FUNED558* | Minas Gerais | Belo Horizonte | Clinical Sample | Swab | 44 | Male | 2024-09-27 |
| 3<sup>#</sup> | FUNED569* | Minas Gerais | Belo Horizonte | Clinical Sample | Swab | 38 | Male | 2024-10-03 |
| 4<sup>#</sup> | FUNED575* | Minas Gerais | Belo Horizonte | Clinical Sample | Swab | 58 | Male | 2024-10-03 |
| 3<sup>#</sup> | FUNED593* | Minas Gerais | Belo Horizonte | Clinical Sample | Swab | 38 | Male | 2024-10-04 |
| 4<sup>#</sup> | FUNED596* | Minas Gerais | Belo Horizonte | Clinical Sample | Swab | 58 | Male | 2024-10-04 |
| - | FUNED726* | Minas Gerais | Belo Horizonte | Environmental | Hospital bed (ICU-21) | - | - | 2024-10-11 |
| - | FUNED727* | Minas Gerais | Belo Horizonte | Environmental | Cables (ICU-21) | - | - | 2024-10-11 |

\*All samples in this table tested positive for *C. auris* using both MALDI-TOF and qPCR assays.
<sup>#</sup>Patients had samples collected twice on different days.

Samples were transported to Lacen-MG in tubes containing Sabouraud Dextrose Broth, processed, and plated onto *Candida* Plus chromogenic agar (Kasvi), followed by incubation for 72 hours at 37°C. Colonies suspected to be *C. auris* were identified by Matrix-Assisted Laser Desorption Ionization Time-of-Flight (MALDI-TOF) mass spectrometry (VITEK® MS – BIOMÉRIEUX).

To confirm the identification, DNA from *C. auris* colonies was extracted and subjected to quantitative PCR (qPCR), following the Centers for Disease Control and Prevention (CDC) protocol (9). For fungal genomic surveillance at Lacen-MG, the DNA from the isolates underwent Whole Genome Sequencing (WGS) using the MiSeq (Illumina) and Ion Torrent PGM (Thermofisher) platforms. Library preparation for Illumina sequencing utilized the *Illumina DNA Prep* and the *600-cycle MiSeq Reagent Kit v3* (Illumina). Sequencing on the Ion Torrent PGM employed the *NEBNext Fast DNA Library Prep kit* (NEB) and the 318 chip (Thermofisher).

Raw sequencing data from the eight *C. auris* isolates were assessed for quality using FastQC (v0.12.1 – github.com/s-andrews/FastQC) and trimmed with Trimmomatic (v0.39 – github.com/usadellab/Trimmomatic). Genome assembly was performed with SPAdes (v4.0.0)(10). Misassemblies were corrected, and contigs were scaffolded using RagTag (v2.1.0 – github.com/malonge/RagTag) with a reference *C. auris* genome (GCA_041381755.1) obtained from GenBank (NCBI). Assemblies were further polished with Pilon (v1.24). The completeness and quality of the assembled genomes were evaluated using QUAST (v5.2.0 – github.com/ablab/quast) and BUSCO (v5.4.7 – github.com/metashot/busco).

To construct a phylogenetic tree, the BUSCO phylogenomics pipeline was used to align 996 single-copy orthologous proteins from 144 *C. auris* genomes (Clades I-VI) available in GenBank (NCBI), along with the eight genomes from this study. IQ-TREE (v2.3.6) was then employed to determine the optimal evolutionary model and construct the phylogenetic tree. Additionally, Parsnp (v1.2 – github.com/marbl/parsnp) was used for core genome alignment, SNP detection, and generating a phylogenetic tree based on the 152 *C. auris* genomes.

To investigate antifungal resistance, the eight genomes generated in this study were analyzed for mutations in key genes associated with resistance: *ERG11* (encoding lanosterol 14 α-demethylase), *ERG6* (encoding sterol 24-C-methyltransferase), *FKS1* (encoding 1,3-beta-glucan synthase), *TAC1B* (encoding a zinc-cluster transcription factor), and *ERG2* (encoding C-8 sterol isomerase) (3,7,11). Know missense mutations linked to antifungal resistance were identified using Blastn (NCBI) to locate target regions, MAFFT (v7.490) for sequence alignment, and AliView for alignment inspection. Detected mutations were validated by mapping sequencing reads to the corresponding gene regions.

*C. auris* was first detected in Brazil in December 2020 (Figure 1a). Since then, it has been reported in four states, with the most recent cases (n=4) identified in Belo Horizonte, Minas Gerais, in September 2024 (Table 1). Eight samples sent to Lacen-MG were confirmed as *C. auris* through MALDI-TOF and qPCR. Epidemiological data (Figure 1b) suggest that the first case in Minas Gerais may have been introduced by a patient hospitalized in Santa Marta, Colombia, and later transferred to Belo Horizonte following an accident. Notably, Patients 1 and 2 occupied the same ICU bed (bed 21) at different times, indicating a potential route for hospital transmission.

**Figure 1.**
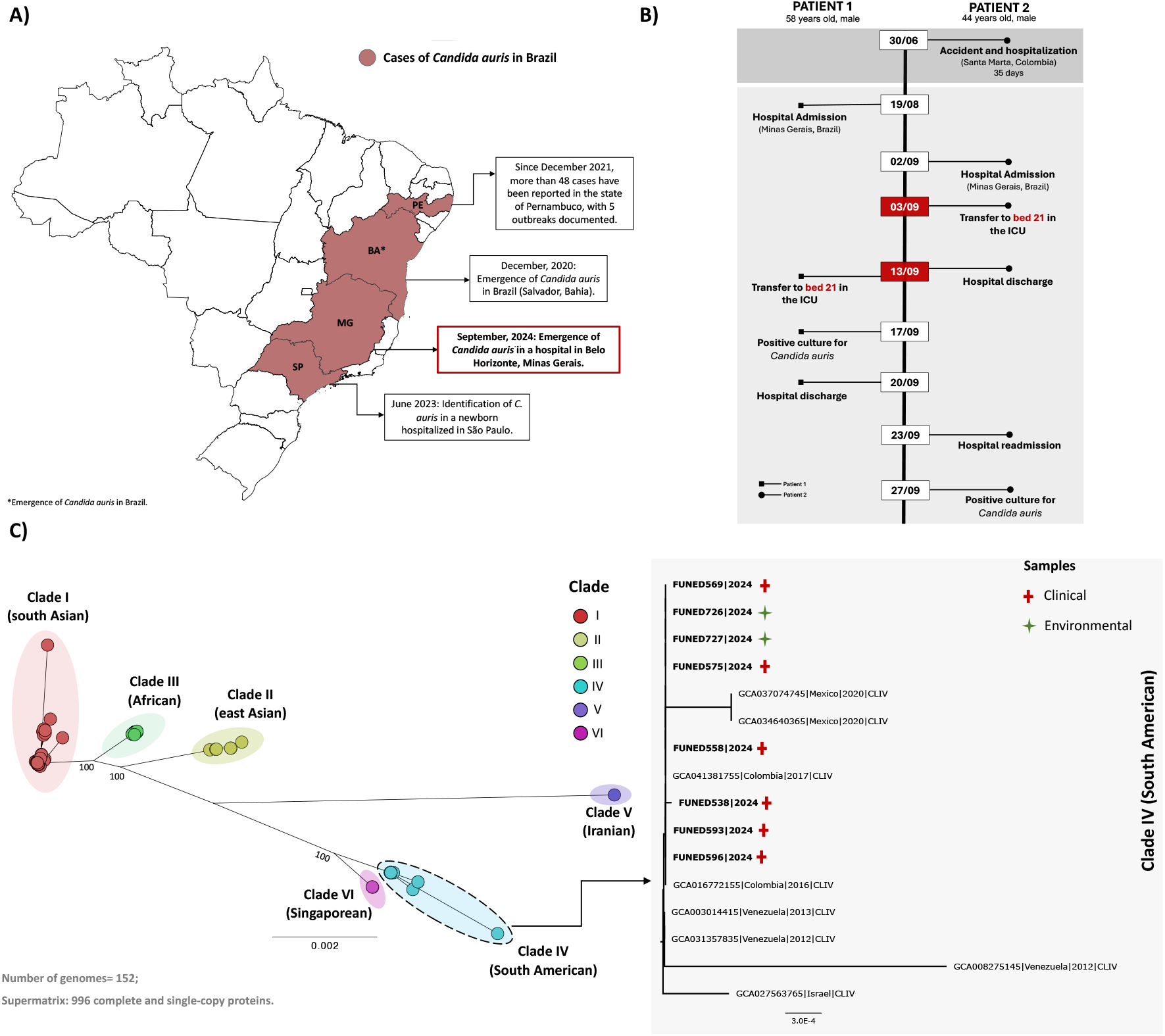
Overview of the emergence of *Candida auris* in Belo Horizonte, Minas Gerais, Brazil. A) Map of Brazil showing *C. auris* detections and outbreaks. B) Timeline of the first *C. auris* cases identified in a hospital in Minas Gerais. Patient 1 was admitted on August 19, 2024, for an aortic dissection. On September 13, the patient was transferred to ICU bed 21. Three days later, they were discharged from the ICU, and on September 17, a urine culture tested positive for *C. auris*. Patient 2 suffered an accident on June 30, 2024, in Santa Marta, Colombia, with two hospitalizations totaling 35 days. After returning to Brazil, they were admitted on September 2 and underwent surgery on September 3, requiring ICU bed 21. C) Phylogenetic tree showing that *C. auris* isolates from Belo Horizonte cluster within Clade-IV, based on 996 concatenated proteins from 152 genomes (8 from this study plus 144 from GenBank – Supplementary Material). The tree was generated using the JTTDCMut+F+R3 model and IQ-TREE with 1,000 bootstrap replicates.

Next-generation sequencing (NGS) provided complete genomes for clinical and environmental isolates, with genome lengths ranging from 12,228,794 to 12,342,789 bp across seven chromosomes, including a complete mitochondrial genome (Table 2). Phylogenomic analysis revealed that all isolates from Minas Gerais belong to Clade-IV, clustering with *C. auris* genomes from Colombia, Mexico, Venezuela, and Israel (Figure 1c). Currently, no Brazilian Clade-IV genomes are available in GenBank (NCBI).

**Table 2.** Genomic metrics of *C. auris* genomes from isolates collected in Belo Horizonte, Minas Gerais, Brazil.

| Sample ID | Reads sequenced | Bases sequenced | Sequencing depth | Genome length (bp) | Number of scaffolds (chromosomes) | N50 | L50 | GC (%) | Complete BUSCOs*(%) |
| --- | --- | --- | --- | --- | --- | --- | --- | --- | --- |
| **FUNED538<sup>#</sup>** | 5,038,765 | 1,020,284,526 | 82 | 12,266,768 | 7 | 2,306,824 | 2 | 45.18 | 98.3 |
| **FUNED558<sup>#</sup>** | 7,076,859 | 1,300,586,893 | 106 | 12,228,794 | 7 | 2,320,025 | 2 | 45.12 | 98.5 |
| **FUNED569<sup>#</sup>** | 3,404,286 | 652,662,068 | 53 | 12,342,789 | 7 | 2,317,092 | 2 | 45.13 | 99.0 |
| **FUNED575<sup>#</sup>** | 5,493,388 | 1,025,455,581 | 83 | 12,332,892 | 7 | 2,316,038 | 2 | 45.12 | 99.0 |
| **FUNED593<sup>#</sup>** | 4,296,743 | 815,830,067 | 66 | 12,279,414 | 7 | 2,307,669 | 2 | 45.12 | 99.0 |
| **FUNED596<sup>#</sup>** | 4,563,919 | 873,120,734 | 70 | 12,327,528 | 7 | 2,315,396 | 2 | 45.12 | 99.0 |
| **FUNED726<sup>#</sup>** | 10,059,594 | 1,840,102,376 | 149 | 12,326,563 | 7 | 2,316,074 | 2 | 45.11 | 99.0 |
| **FUNED727<sup>#</sup>** | 8,052,864 | 1,520,151,176 | 123 | 12,336,396 | 7 | 2,315,903 | 2 | 45.12 | 99.0 |

<sup>#</sup>All genomes will be deposited and made publicly available under BioProject ID: PRJNA1194488 (NCBI).
\*BUSCO analysis based on the presence of complete single-copy orthologous genes.

SNP-based analysis demonstrated that all genomes clustered within their respective clades in the SNP phylogenetic tree and PCA plot (Figures 2a and 2b). Consistent with the protein-based phylogeny (Figure 1c), the SNP tree (Figure 2c) confirmed that the Belo Horizonte genomes are part of Clade-IV clustering closely with two Colombian genomes, supporting a potential introduction from Colombia. Furthermore, the genome from Patient 2, who had been previously hospitalized in Colombia (FUNED558) appeared as an early branch, suggesting it may represent first case in Belo Horizonte.

**Figure 2.**
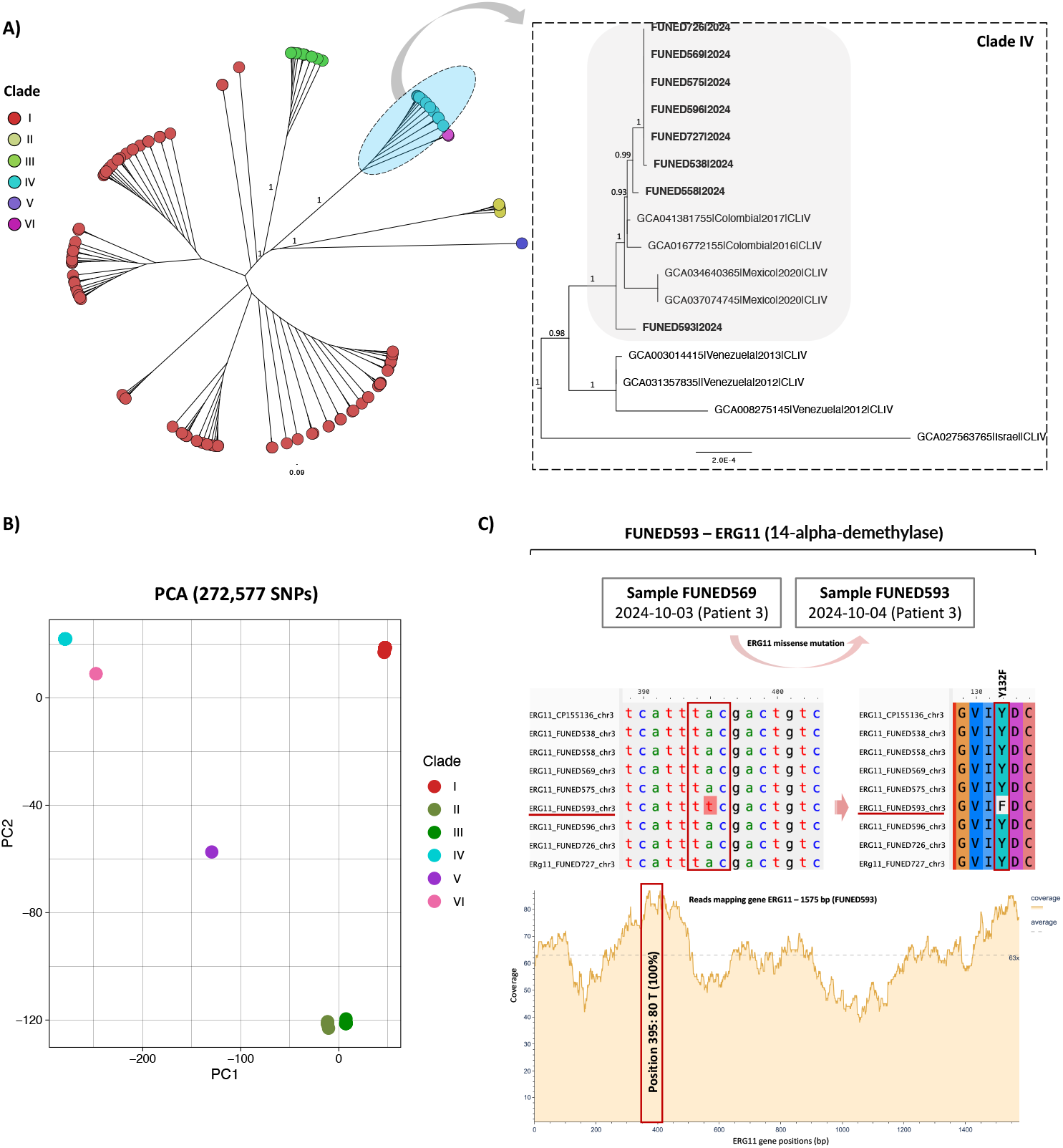
*Candida auris* genome analysis based on Single Nucleotide Polymorphism (SNP) detection and antifungal resistance mutations. A) Phylogenetic tree based on core genome alignment and SNP detection using Parsnp. Support values for major clades are shown. B) Principal Component Analysis (PCA) plot derived from a matrix of 272,577 SNPs from 152 genomes, generated using Parsnp, Gingr (VCF extraction), and R libraries (vcfR, dplyr, ggplot2, readxl, tidyr). C) Analysis of the ERG11 gene (encoding 14-alpha-demethylase) showing a A-to-T nucleotide mutation at position 395 in the FUNED593 genome. This results in an amino acid substitution from tyrosine (Y) to phenylalanine (F) at position 132. The ERG11 sequences from this study were aligned with chromosome 3 (CP155136.1) of the GCA_041381755.1 genome. Mapping reads to ERG11 gene shows a sequencing depth of 80X at the mutation site in the FUNED593 genome.

Notably, in Colombia, *C. auris* was retrospectively identified as circulating since 2012 in Santa Marta, the city of origin for Patient 2 (12). Additionally, all genomes analyzed in this study exhibit amino acid substitutions in the *ERG11* gene (K177R, N335S, E343D), which is characteristic of Colombian isolates (13).

The analysis of antifungal resistance in the FUNED593 genome (Patient 3) identified a missense mutation in *ERG11*, a gene critical for ergosterol biosynthesis and the target of azole antifungals. This mutation resulted in a Y132F amino acid substitution in *Erg11p*. This alteration is well-documented as a mechanism conferring fluconazole resistance in *C. auris* and other *Candida* species (3,4,11,14)(Figure 2c). Interestingly, this mutation was absent in the sample collected from the same patient the previous day. This is the first identification of this mutation in Brazilian *C. auris* isolates, underscoring the critical need for ongoing genomic characterization to monitor and further understand the mechanisms underlying such mutations.

## Conclusions

These findings confirm the first detection of *C. auris* in Belo Horizonte, likely introduced from Colombia, with evidence of hospital transmission. Phylogenomic analyses identified critical genomic traits and emerging mutations that can confer antifungal resistance. The study emphasizes the importance of genomic surveillance for tracking pathogen spread and resistance mutations, guiding public health responses. It also highlights that *C. auris* colonization alone can rapidly select for resistance mutations, reinforcing the need for rigorous hospital surveillance and containment measures to prevent further spread.

## Acknowledgments

We thank the Ezequiel Dias Foundation (Funed) for providing infrastructure and funding for this Laboratory and Genomic Surveillance project. We also thank the Laboratory of Molecular and Computational Biology of Fungi (LBMCF) at the Federal University of Minas Gerais (UFMG) for providing high-performance computers for bioinformatics analyses. We are grateful to the Secretaria de Estado de Saúde de Minas Gerais (SES-MG) for their support in laboratory surveillance efforts. L.M.R. Tomé and N.R. Guimarães received scholarships (BDCTI-I) from the Fundação de Amparo à Pesquisa do Estado de Minas Gerais (FAPEMIG) through project RED-00234-23. T.E.R. Adelino and S.H.S.P. Pedroso are supported by the Conselho Nacional de Desenvolvimento Científico e Tecnológico (CNPq) under process numbers 153597/2024 and 172892/2023-6, respectively. M. Giovanetti received funding from the PON “Ricerca e Innovazione” 2014–2020 program. This study was also financed by National Institutes of Health (grant no. U01 AI151698) for the United World Arbovirus Research Network (UWARN).

## About the author

Luiz Marcelo Ribeiro Tomé holds a Ph.D. in microbiology with a focus on mycology and bioinformatics. He completed his postdoctoral training in bioinformatics at UFMG. He is currently a postdoctoral researcher at Fiocruz Minas and Funed, specializing in genomic surveillance of emerging and re-emerging pathogens, emphasizing bioinformatics analyses.

## Notes

### Competing Interest Statement

The authors have declared no competing interest.

